# Two-species community design of Lactic Acid Bacteria for optimal production of Lactate

**DOI:** 10.1101/2020.10.24.353805

**Authors:** Maziya Ibrahim, Karthik Raman

## Abstract

Microbial communities that metabolise pentose and hexose sugars are useful in producing high-value chemicals, as this can result in the effective conversion of raw materials to the product, a reduction in the production cost, and increased yield. Here, we present a computational approach called CAMP (Co-culture/Community Analyses for Metabolite Production) that simulates and identifies appropriate communities to produce a metabolite of interest. To demonstrate this approach, we focus on optimal production of lactate from various Lactic Acid Bacteria. We used genome-scale metabolic models (GSMMs) belonging to *Lactobacillus, Leuconostoc*, and *Pediococcus* species from the Virtual Metabolic Human (VMH; https://vmh.life/) resource and well-curated GSMMs of *L. plantarum* WCSF1 and *L. reuteri* JCM 1112. We studied 1176 two-species communities using a constraint-based modelling method for steady-state flux-balance analysis of communities. Flux variability analysis was used to detect the maximum lactate flux in a community. Using glucose or xylose as substrates separately or in combination resulted in either parasitism, amensalism, or mutualism being the dominant interaction behaviour in the communities. Interaction behaviour between members of the community was deduced based on variations in the predicted growth rates of monocultures and co-cultures. Acetaldehyde, ethanol, NH_4_^+^, among other metabolites, were found to be cross-fed between community members. *L. plantarum* WCSF1 was a member of communities with high lactate yields. *In silico* community optimisation strategies to predict reaction knock-outs for improving lactate flux were implemented. Reaction knock-outs of acetate kinase, phosphate acetyltransferase, and fumarate reductase in the communities were found to enhance lactate production.

**Importance:** Understanding compatibility and interactions based on growth between the members of a microbial community is imperative to exploit these communities for biotechnological applications. Towards this goal, here, we introduce a computational analysis framework that evaluates all possible two-species communities generated from a given set of microbial species on single or multiple substrates to achieve optimal production of a target metabolite. As a case study, we analysed communities of Lactic Acid Bacteria to produce lactate. Lactate is a platform chemical produced experimentally from lignocellulosic biomass, which constitutes pentoses and hexoses, such as xylose and glucose. Metabolic engineering strategies, such as reaction knock-outs that can improve product flux while retaining the community’s viability are identified using *in silico* optimisation methods. Our approach can guide in the selection of most promising communities for experimental testing and validation to produce valuable bio-based chemicals.

## Introduction

In recent years, novel methods for synthesising valuable chemicals include the use of co-cultures or microbial communities, where two or more microbial populations are cultured together to derive optimum output of the product (1). In nature, microbes exist in communities, and the use of natural or engineered consortia have advantages over single strains. One of the critical features of a consortium is the division of labour or sharing of metabolic burden between the species. The product of one engineered strain is transported to another microbe, where it can be further metabolised to the final desired metabolite. Co-cultures allow a symbiotic relationship between strains for the utilization of multiple substrates and removal of inhibitory by-products. Some challenges in co-culture studies include compatibility between the strains concerning their growth conditions such as temperature, pH, and media (2).

Computational modelling of co-cultures is feasible with the use of genome-scale metabolic models (GSMMs). GSMMs of micro-organisms computationally describe the metabolism of an organism through the gene-protein-reaction associations. Progress in the reconstructions of GSMMs has allowed a wide variety of metabolic studies by generating model-driven hypotheses and context-specific simulations by the integration of various omics and kinetic data. GSMMs have been used to predict targets for gene manipulation either through knock-out or up- and downregulation, which has resulted in improved production of industrially relevant chemicals from micro-organisms (3). In an *E. coli* strain (XB201T) producing 0.55 g/L of D-phenyl lactic acid, knock-outs of *tyrB* and *aspC* genes that were identified as potential knock-out candidates from in silico analysis enhanced the production to 1.62 g/L (3).

The use of constraint-based modelling approaches with microbial community models is also underway to study metabolic interactions between the species (4–6). In the current study, we present a constraint-based modelling approach called CAMP (Co-culture/Community Analyses for Metabolite Production) which evaluates a set of GSMMs to identify suitable two-species communities that can produce a given metabolite. We demonstrate this approach by analysing GSMMs of selected Lactic Acid Bacteria (LAB) to construct two-species communities and examine their potential for optimal production of lactate.

Lactate is an α-hydroxy carboxylic acid that is chemically reactive and is synthesised to various intermediates such as acrylic acid, 1,2-propanediol, and lactide. Lactide is the building block for producing polylactic acid (PLA) (7). PLA is a biodegradable biopolymer that finds applications in the biomedical industry to manufacture stents, surgical sutures, soft-tissue implants, etc. (8). Lactic acid is also used in the food industry as an acidulant, a preservative, and an emulsifier (7). The D-isomer is considered harmful to humans in high doses. It can cause acidosis or de-calcification; hence, the L-isomer of lactate is preferred in the food and pharmaceutical industry (9).

Microbial fermentation is an effective route to produce lactate, as optically pure D- or L-lactate can be produced based on the selection of appropriate micro-organisms. LAB can be classified as either homofermentative or heterofermentative, depending on the metabolism of hexoses and pentoses, and the production of end products. In homofermentative cases, the sugars are metabolised via the Embden-Meyerhof-Parnas (EMP) pathway, whereas in the heterofermentative case, the phosphoketolase pathway is active (10).

In *Lactobacillus* co-cultures of *L. brevis* and *L. plantarum* with glucose and xylose as substrates and NaOH treated corn stover, high lactate yields of 0.8 g/g were obtained, which is more significant than in monocultures of the same species (11). *L. rhamnosus* and *L. brevis* were also used in co-culture, and a lactate productivity of 0.7 gL^-1^h^-1^ was obtained (12). Co-culture of *L. pentosus* and genetically engineered *Enterococcus faecalis* produced 3.68 gL^-1^h^-1^ of lactate (1). A consortium of cellulolytic fungus *Trichoderma reesei* and *L. pentosus* fermented on whole-slurry pre-treated beech wood led to the production of 19.8 g/L of lactic acid. *L. pentosus* consumed cellobiose, avoiding inhibition of *T. reesei* cellulase activity, and acetic acid produced from *L. pentosus* was utilised as a carbon source by the fungus (13). GSMMs of various LAB such as *Lactobacillus reuteri, Leuconostoc mesenteroides, Lactobacillus plantarum, Lactobacillus casei, Lactococcus lactis*, and *Streptococcus thermophilus* have been published (14).

We used the CAMP approach to predict growth rates of LAB species in monoculture and co-culture. We categorised the interactions in LAB communities based on the changes in predicted growth rates, either unidirectional such as commensalism, amensalism, and neutralism, or bi-directional such as mutualism and competition. We analysed the effects of single and multiple nutrient substrates on interaction types between communities. We examined the metabolites that are exchanged between the species of a community. We predicted reaction knock-outs in LAB communities that would improve lactate flux. Overall, our strategy is generic, and it can be applied to identify communities to produce specific metabolites of interest. We postulate that this analysis strategy will benefit metabolic engineering applications that involve microbial communities.

## Results

In this section, we present a brief overview of the CAMP approach, followed by its application to identify the most promising co-cultures to produce lactate.

### Overview of CAMP (Co-culture/Community Analyses for Metabolite Production)

Fig. 1 gives an outline of the CAMP workflow. The steps include 1) Retrieval of microbial GSMMs from databases such as VMH. Each of these GSMMs is simulated in three different nutrient conditions (See Materials ad Methods). Predicted growth rates and product flux are obtained using flux balance analysis (FBA) and flux variability analysis (FVA). The product yield is computed as the maximum product flux obtained per unit flux of substrate uptake. 2) Two-species communities are created using SteadyCom (6). Community models are also simulated in three nutrient conditions. FBA and FVA are used to predict community growth rates and product yield in the community. Monoculture and co-culture growth rates are compared to identify an increase or decrease in growth when an organism is simulated in the presence of another. 3) Expected product yield in a community is compared to the observed product yield. Details on calculation of product yield can be found in Materials and Methods. Communities which have a 10-fold increase in product yield are regarded as candidate communities for optimal production of the target metabolite. Communities are assessed for their relative abundances, type of interaction behaviour observed and the cross-fed metabolites. 4) *In silico* community optimisation is performed using FSEOF (15), which enables to shortlist potential reaction knock-outs that will increase product flux in the community. Reaction knock-outs can be from either species in the community.

**Fig. 1.**
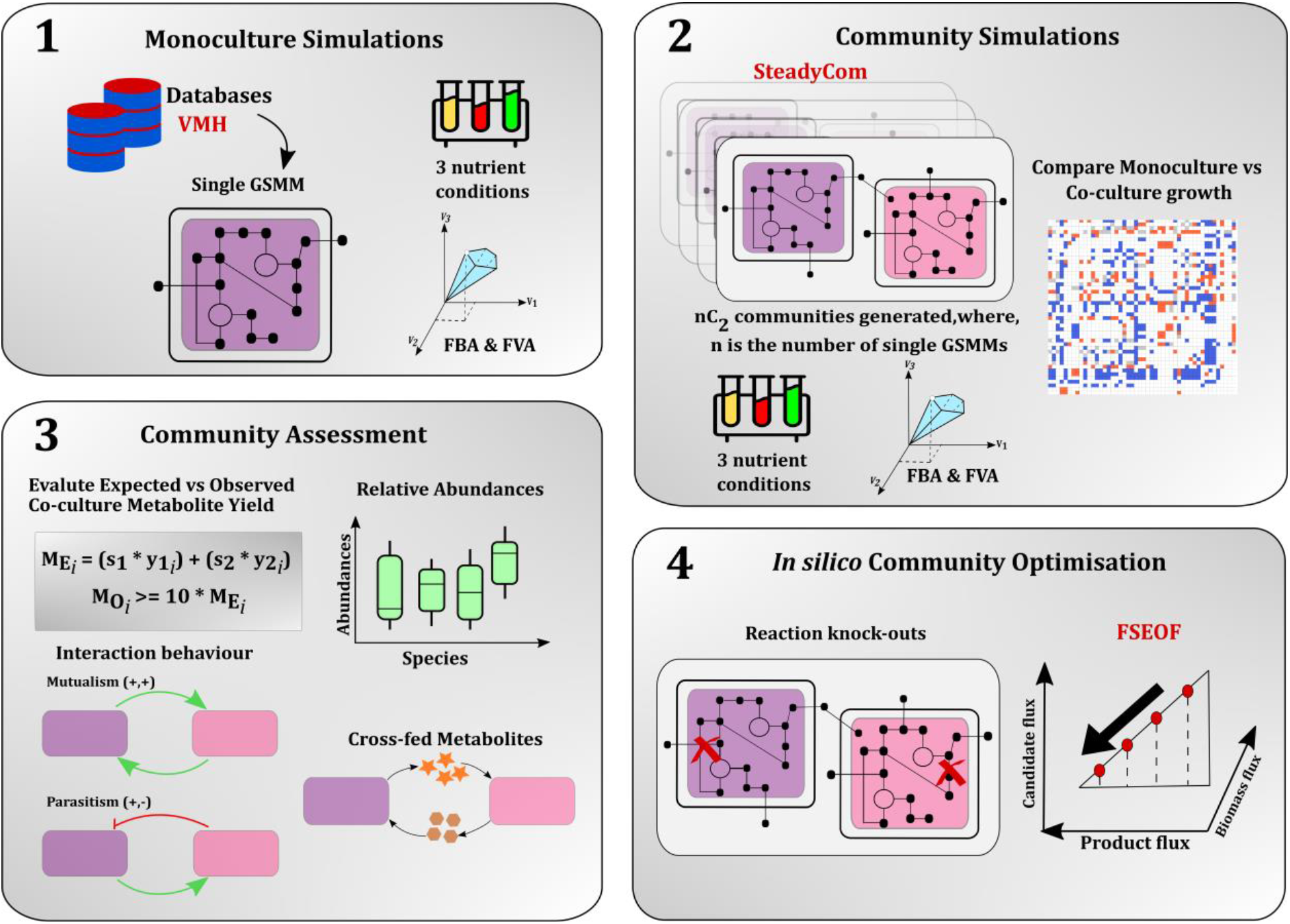
Outline of CAMP (Co-culture/Community Analyses for Metabolite Production)

### Growth phenotypes of LAB in monoculture

For all 49 GSMMs, their predicted growth rates in monoculture with glucose and xylose as major carbon sources were computed for the three different nutrient conditions — minimal nutrient, excess nutrient, and community-specific nutrient condition (see Materials and Methods). The maximal lactate fluxes of each model in all three conditions were also computed. Supplementary Table S1 details the growth rates of each LAB species in the different nutrient conditions. It was observed that for all models, the active reactions that had a non-zero flux belonged to the central carbon metabolism, such as Embden-Meyerhof-Parnas (EMP) pathway, pentose phosphate pathway (PPP), and the pentose phosphoketolase (PPK) pathway (16) as seen in Fig. 2. Histogram distribution of predicted monoculture growth rates (Supplementary Fig. S1) under the three nutrient conditions shows that many species have similar growth rates in all conditions within the range of 0.01 to 0.1 (h^-1^). The highest growth rates (> 0.3 h^-1^) are observed in the community-specific and excess nutrient conditions.

**Fig. 2:**
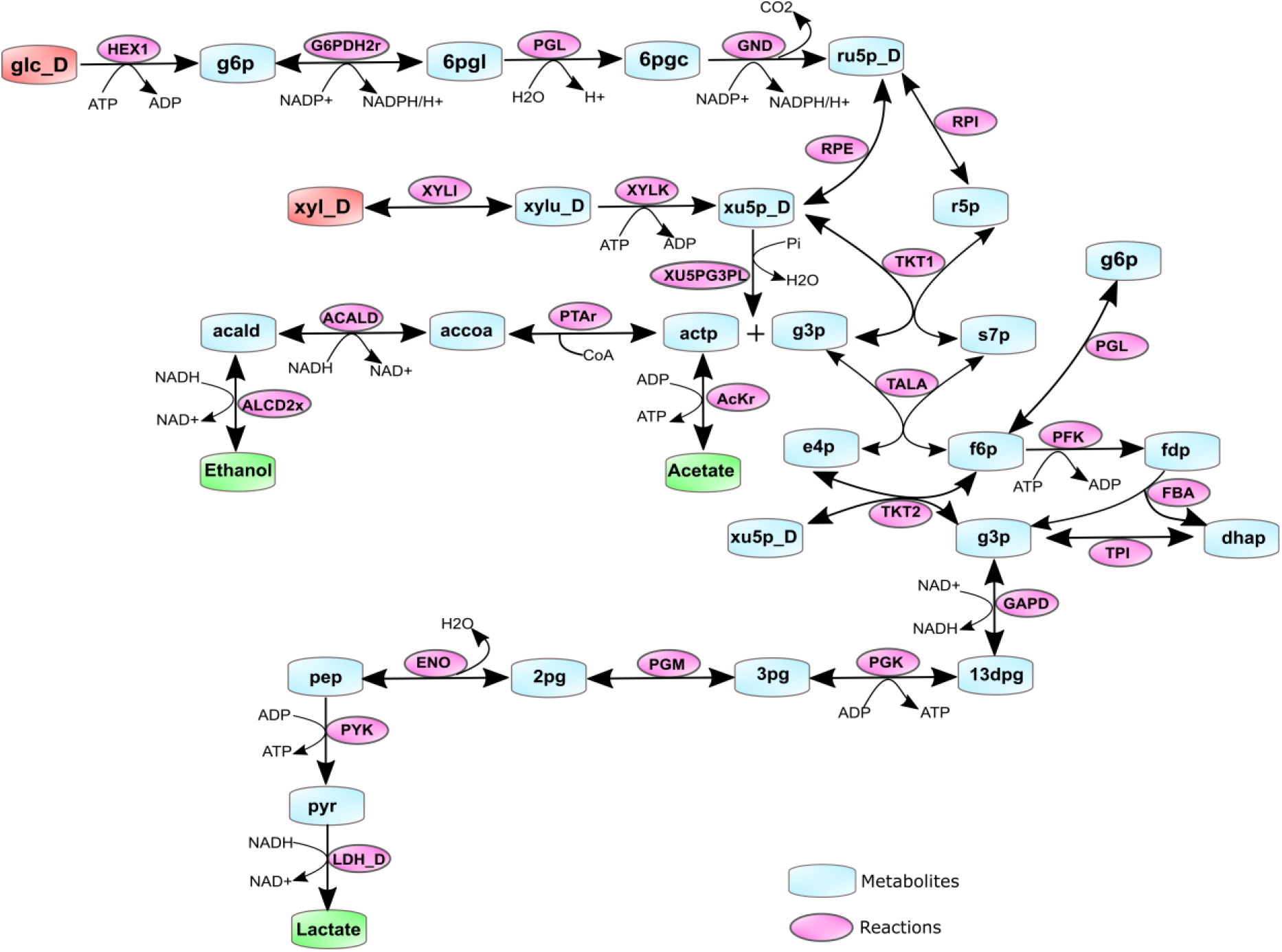
Active pathway reactions with non-zero fluxes in the LAB models when grown in monoculture and co-culture. Glucose and xylose (shaded red) are the primary substrates that are metabolised to the end-products lactate, acetate, and ethanol (shaded green). Metabolite and reaction notations and reaction directionalities are denoted as seen in the LAB GSMMs.

### Significant change in monoculture vs. co-culture growth rates helps segregate communities into six categories

A difference of 10% in predicted growth rates of the microbes in monoculture versus co-culture has been previously established to be significant (17). Based on these comparisons, viable LAB communities from each nutrient condition were binned into categories as follows: Amensal communities, i.e., one microbe grows slower in the paired simulation while the other microbe’s growth rate is unaffected. Competitive communities, i.e., both microbes’ growth, is slower than their monoculture rates. Parasitic communities, i.e., one microbe grows faster in the paired simulation while the other microbe grows slower. Neutral communities, i.e., neither microbes’ growth rate was affected upon being paired with the other. Commensal communities, i.e., one microbe, has an increase in growth rate while the other remains unaffected. Lastly, mutualistic communities where both microbes in the pair show an increase in the growth rates compared to their monoculture rates. Fig. 3 depicts the interaction behaviour in communities when each microbe influences the growth of the other, either positively or negatively.

**Fig. 3.**
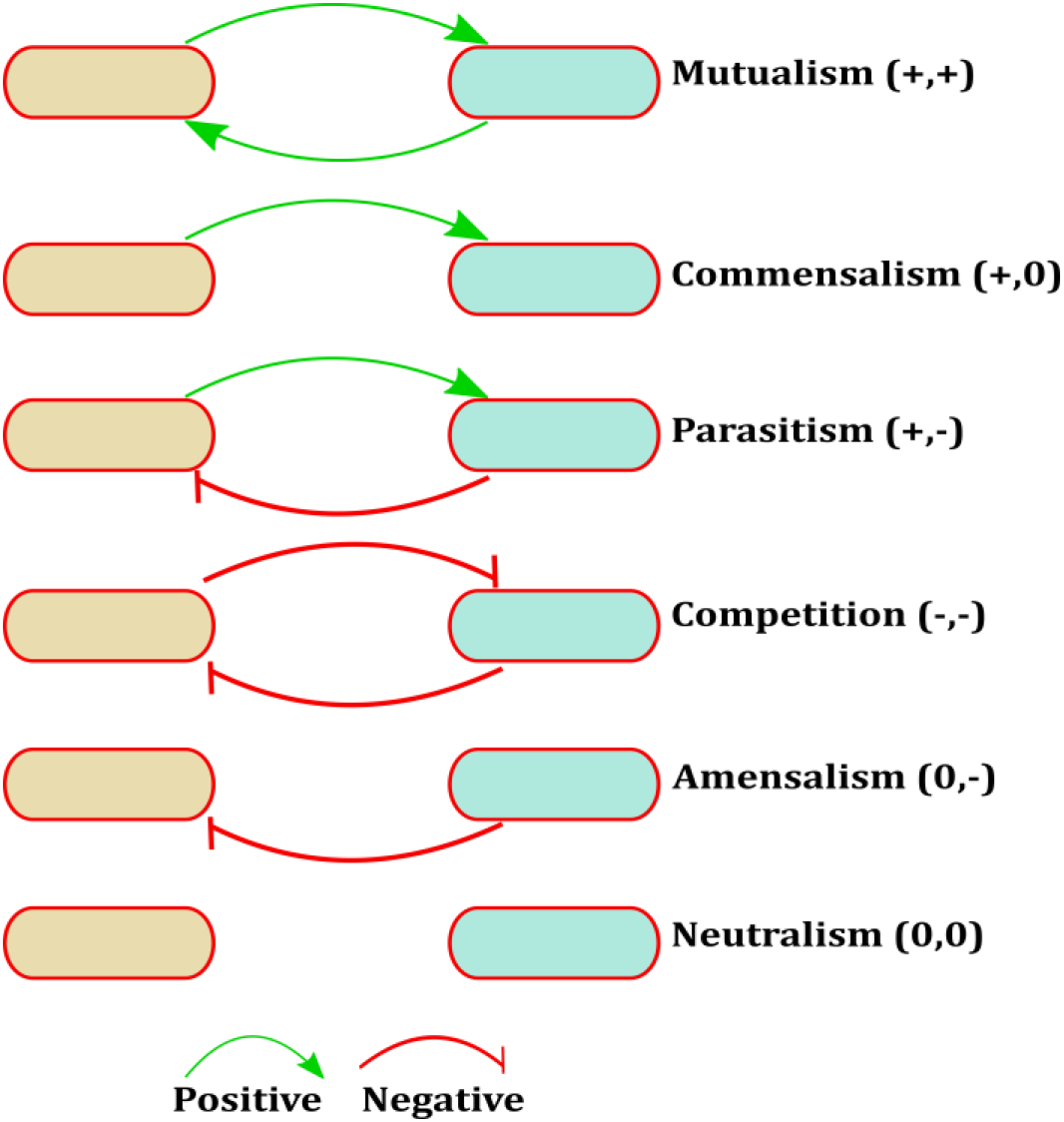
Different interaction types possible between the two-species communities. A positive or negative effect on the growth of the species defines each interaction type.

In community-specific nutrient conditions, 354 viable pairs out of 1176 were identified, as seen in Fig. 4. Parasitism was the ‘favoured’ interaction type, with 235 pairs out of 354 displaying parasitic behaviour. In minimal nutrient conditions, there were 492 viable pairs. Again, parasitism was dominant in this group, with 224 out of 492 pairs exhibiting parasitism. In contrast, in the excess nutrient condition, from among 338 viable pairs, 215 pairs had amensal behaviour. Parasitism, mutualism, and commensal pairs were not identified in this group. Heatmaps for the minimal and excess nutrient conditions are provided as supplementary Fig. S2 & Fig. S3. Supplementary Fig. S4, S5 and S6 contain heatmaps that depict the absolute values of the predicted growth rates of each species grown in the presence of 48 other species.

**Fig. 4:**
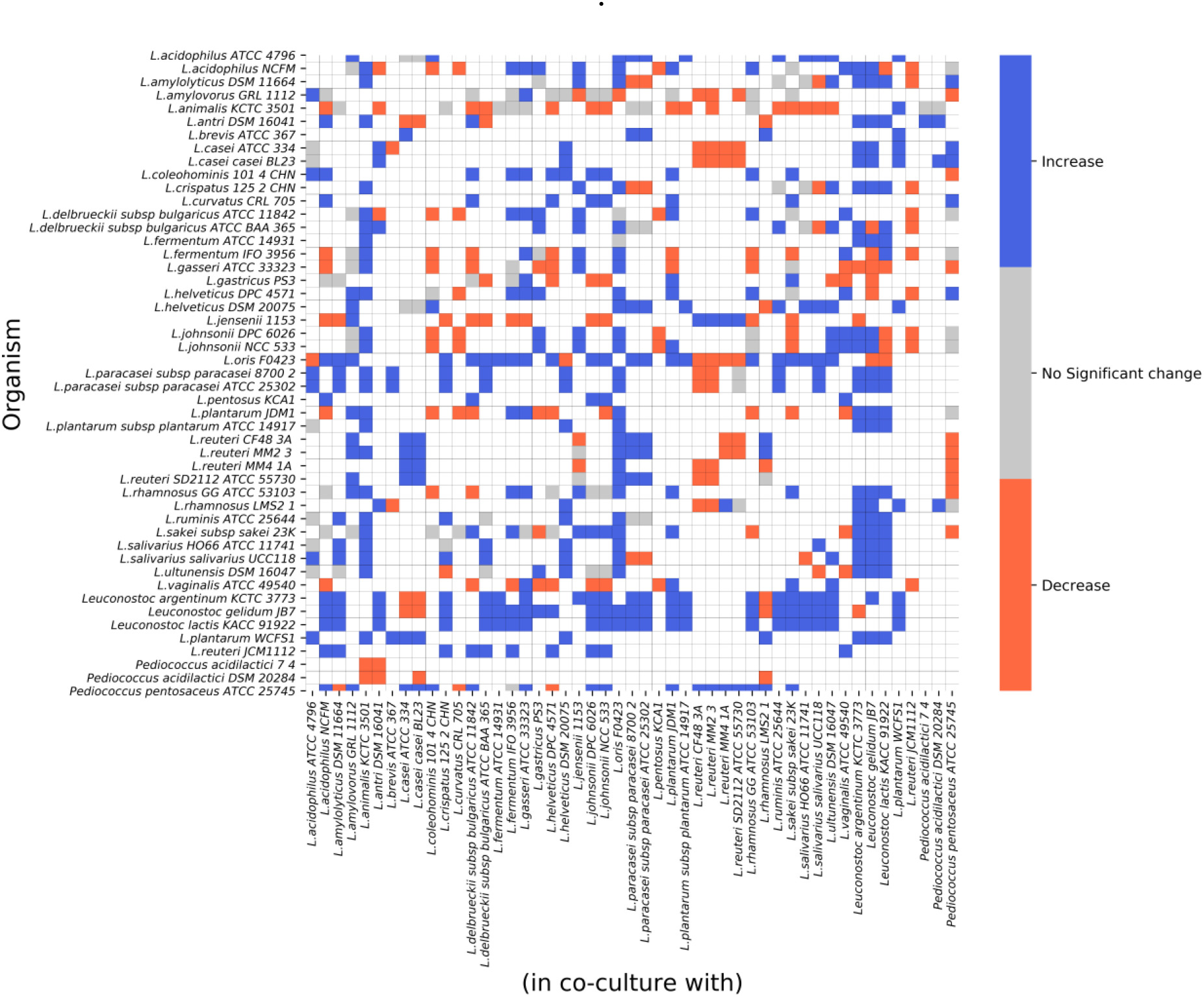
Monoculture vs. co-culture growth rates. The heatmap depicts the change in the growth rate of an organism’s predicted monoculture growth compared to when it is co-cultured with another species under community-specific nutrient condition. A difference greater than 10% of monoculture growth is considered an increase, whereas lesser than 10% of monoculture growth is regarded as a decrease. 822 non-viable pairs and the diagonal, which represents 49 monocultures, are depicted as white squares.

### Occurrences and relative abundance profiles of the LAB species

The frequency of occurrence of each microbe among the viable communities in each nutrient condition was calculated. *L. oris* and *L. animalis* had the highest occurrences among all *Lactobacillus* species. *Leuconostoc* species were also found to rank higher in thenumber of occurrences among the viable set, irrespective of the nutrient condition. Each of these microbes was found in at least 20 pairs or more. *Pediococcus* species formed the least number of pairs in the community-specific nutrient condition. *L. pentosus* KCA1 was found to constitute the least number of viable pairs (less than 10) in all nutrient conditions.

The distribution of predicted relative abundances of each microbe when co-cultured under different nutrient conditions are shown in Fig 5. The abundances were found to vary depending upon the number of viable communities associated with each microbe. Differences were also seen among the nutrient conditions, with most LAB species having a mean abundance of lesser than 0.5 in the excess nutrient condition. *L. oris*, present in many viable communities, had an average abundance of less than 0.25 in the minimal and excess nutrient conditions. In contrast, it had an abundance higher than 0.5 in the community-specific condition. Relative abundances greater than 0.75 were seen among *Leuconostoc* species and some *Lactobacilli* species in the community-specific nutrient condition. This variation in abundance profiles highlights the role of nutrient constraints in driving community behavior.

**Fig. 5.**
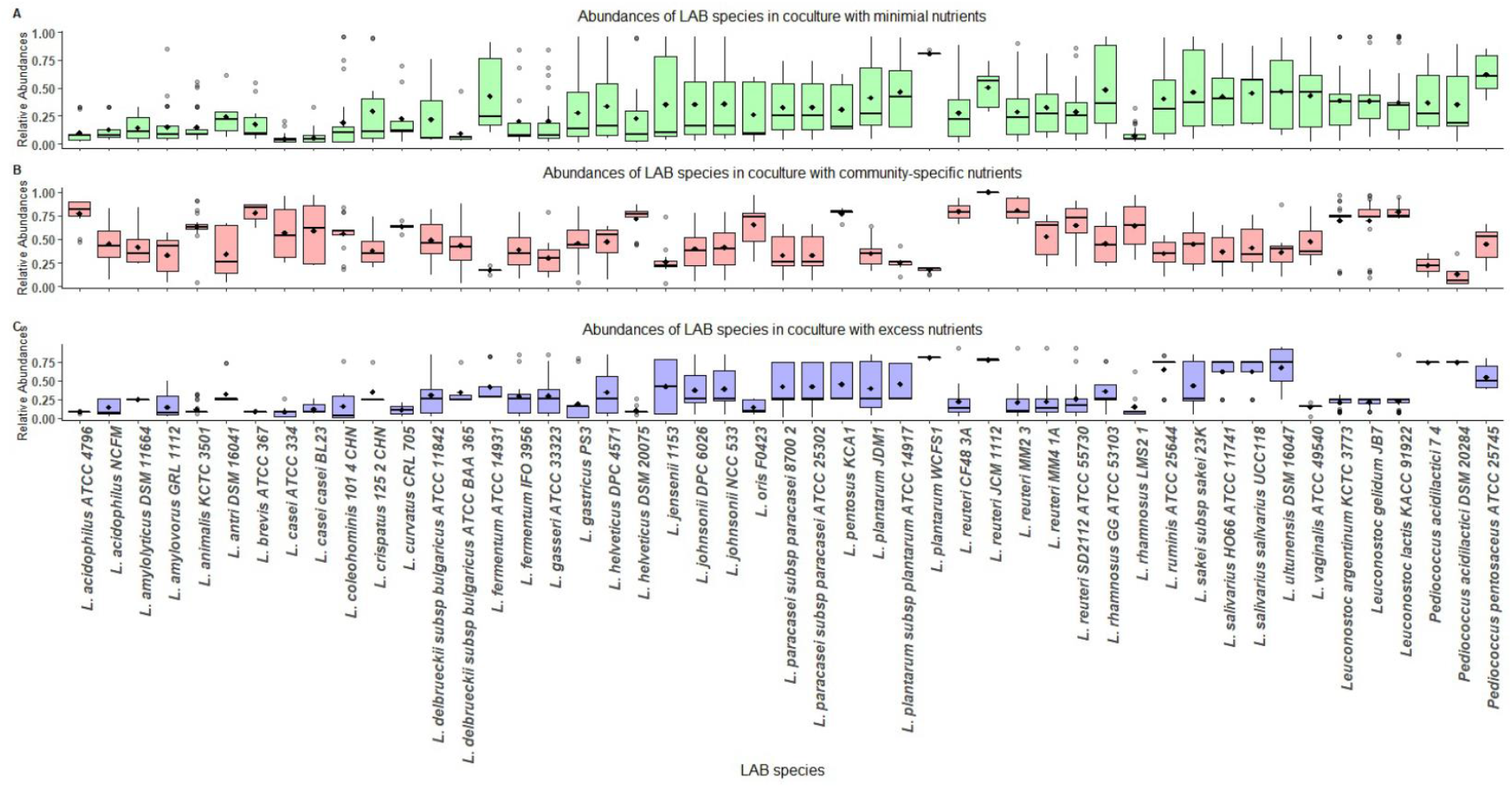
Relative abundance profiles of LAB species in co-culture under different nutrient conditions. (A) minimal nutrient condition (B) community-specific condition (C) excess nutrient condition.

### Dominant interaction behavior differs in communities grown with single and multiple substrates

To examine if the type of interaction detected in a community is dependent on the number of carbon sources utilised, we simulated the community models for growth on glucose and xylose independently. We compared these findings to when both glucose and xylose are provided as substrates to the communities for growth. Fig. 6 highlights the interaction types observed when either glucose or xylose is used as a substrate under different nutrient conditions.

**Fig. 6.**
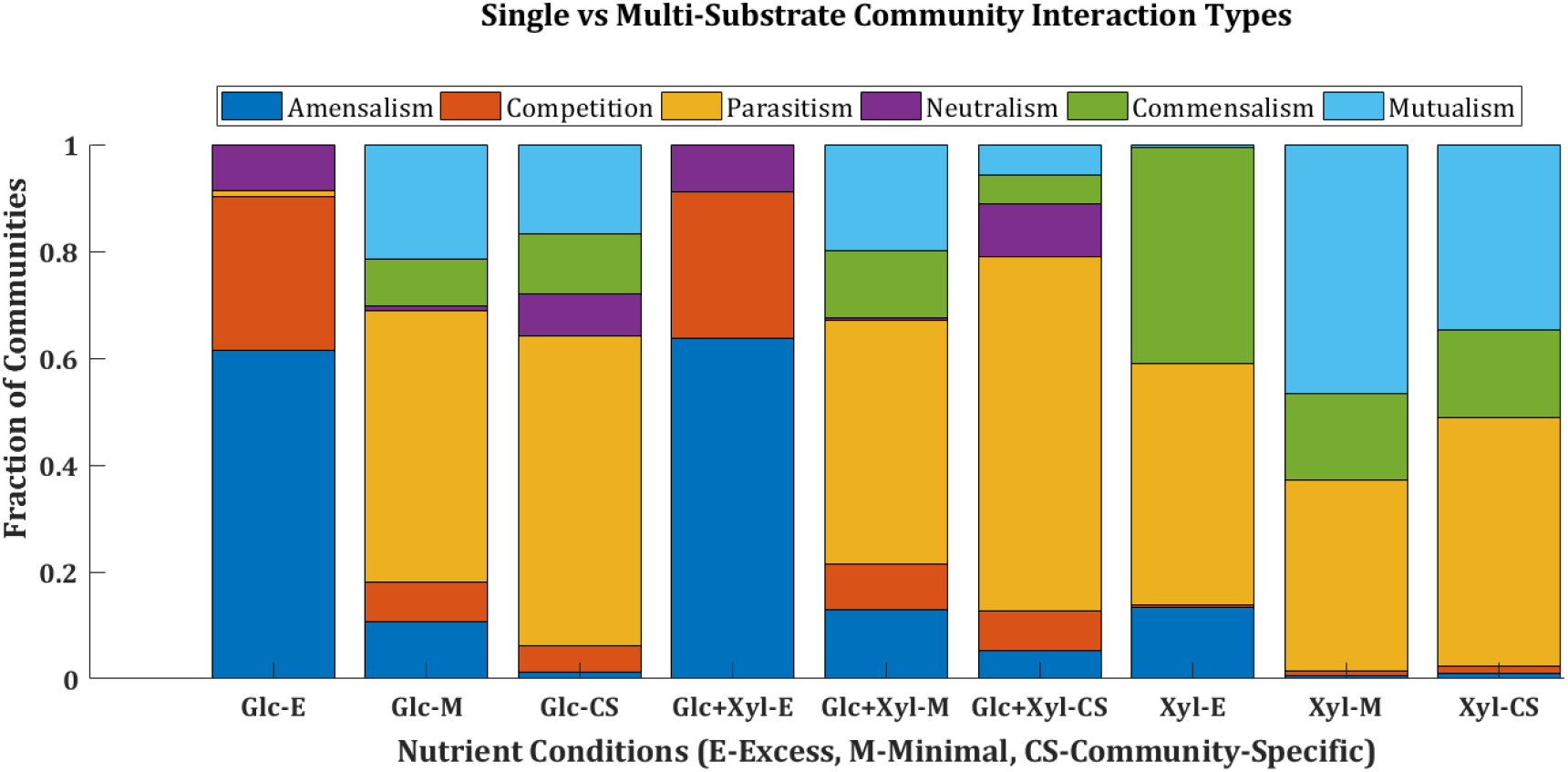
Distribution of the various interaction types between viable pairs in nine different nutrient conditions. The plot shows the fraction of communities with a particular interaction type in each nutrient condition.

Among the 49 LAB models, only 11 models can metabolise xylose as a sole nutrient source. Mutualistic pairs constituted an average of 40% of viable pairs in minimal and community-specific conditions with xylose as substrate. The number of mutualistic pairs in xylose-only conditions indicates the rise of an emergent property in the community. Viable pairs with amensalism behaviour are found to be higher in excess nutrient conditions. Parasitism prevailed in both minimal and community-specific nutrient conditions irrespective of the presence of a single or multi-substrate. As all 49 organisms are capable of metabolising glucose, some competitive behavior is observed primarily in glucose-only excess conditions. Whereas, in xylose-only conditions, competition is almost absent, with only a maximum of three viable pairs exhibiting competition.

### Communities possess positively and negatively correlated cross-fed metabolites

A metabolite was considered cross-fed if it was secreted (i.e., the flux of the exchange reaction for the particular metabolite was positive) into the community compartment (u)by one organism and taken up (i.e., the flux of the exchange reaction of the metabolite was negative) by the other organism in the community. A threshold of 0.01 mmol/gDW/h was used to determine all such cross-fed metabolites for the viable communities in each nutrient condition. Fifty-three unique metabolites that included many amino acids were cross-fed between the LAB communities. This is consistent with other experimental observations where the exchange of amino acids is considered to play a role in community interactions (18, 19). The most widely cross-fed metabolites across all viable communities were acetaldehyde, glycine, H^+^, ethanol, H_2_O, acetate, formate, and NH_4_^+^. Lactate was also found to be cross-fed between 35% of communities across different nutrient conditions. Each community model exchanged varied sets of metabolites depending on the nutrient condition it was simulated in. To check if certain metabolites are always cross-fed simultaneously in a community, the correlation between cross-fed metabolites across the LAB communities was estimated (Fig. 7). In the communityspecific nutrient condition, positively correlated metabolites with a *p*-value significance of less than 0.05 (adjusted by the Benjamini-Hochberg method to control the false discovery rate) were identified to be ethanol and H_2_O, stearic acid and hypoxanthine, and formate and serine. Negatively correlated metabolites were formate and H_2_O, glycerol and acetaldehyde. We checked whether the cross-fed metabolites are specific to any interaction type and found that 24 metabolites are common to all interaction types. They include succinate, malate, formate, ethanol, acetate and some amino acids. The fraction of metabolites cross-fed in cooperative communities with mutualistic, commensal, and neutral interactions are higher than in communities which exhibit parasitic and competitive behaviour. Supplementary Table S2 has the list of cross-fed metabolites in each interaction type.

**Fig. 7.**
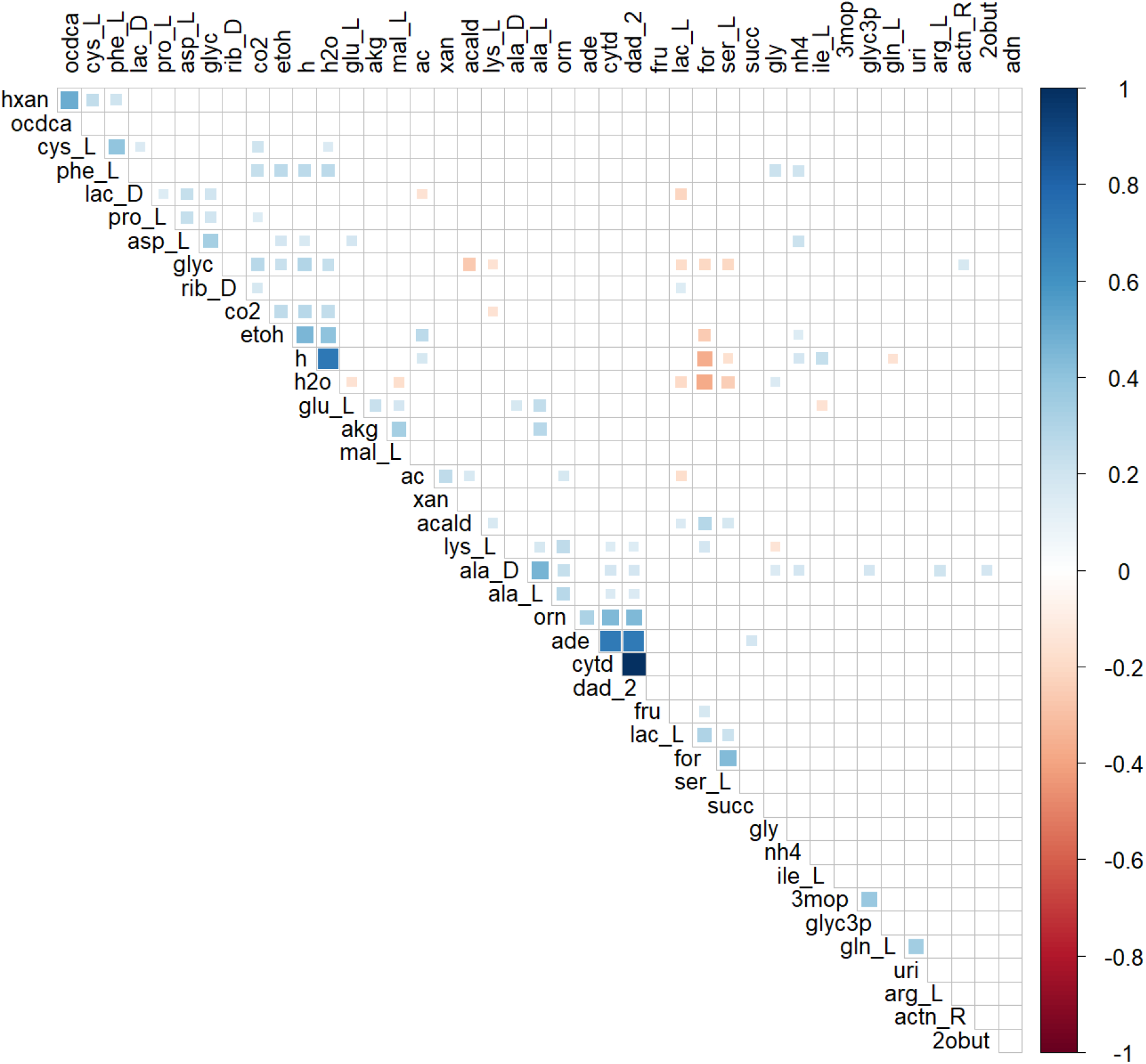
Correlation between the cross-fed metabolites in the community-specific nutrient condition. Positively correlated metabolites are denoted in blue, whereas negatively correlated metabolites are coloured brown. Correlation plots for cross-fed metabolites in the other two nutrient conditions are provided as Supplementary figures, Fig. S7 and Fig. S8

### Evaluating performance of communities based on growth and lactate yield

We evaluated the performance (see Materials and Methods) of the community models in two scenarios. In the first set of simulations, lactate was not allowed to be cross-fed between the community members. In the second case, one organism in the pair is designated as the primary consumer of the substrates glucose and xylose, thereby creating a dependence of the second organism on the first for growth and vice-versa. Community pairs that retained their viability in the two test scenarios were deemed fit for further community strain optimisation strategies. This performance test was carried out in all three nutrient conditions. Forty community pairs were common in two nutrient conditions, community-specific nutrient uptake and minimal nutrient uptake. Seven LAB communities were unique to the excess nutrient condition. Each of these pairs had an observed lactate yield 10-fold higher than the expected lactate yield of the community.

### Glucose fermenters have higher lactate yield than communities where both xylose and glucose is utilised

For grading the community pairs based on both their growth rate and product yield, the biomass, and lactate flux values were normalised (min-max normalization). Upon normalisation, the best pairs were identified. A detailed list of all communities is found in Supplementary Table S3. Each of the top six pairs shared an organism, namely, *L. plantarum* WCFS1, which is coupled with two strains of *L. casei, L. rhamnosus* LMS2, *L. animalis* KCTC 3501, *Leuconostoc argentinum*, and *Leuconostoc lactis*.

Contrary to expectations, in the best-performing pairs, both the organisms are not capable of utilising glucose and xylose together. Only the *Leuconostoc* species can metabolise both glucose and xylose, while the remaining organisms are glucose fermenters. The metabolic distances (Jaccard distances) between the GSMMs in the best-performing pairs were calculated (see Materials and Methods) using reaction lists from each model. The top-ranked pairs had a Jaccard distance of greater than 0.7, indicating that they had less than 30% of their reactions in common, and therefore, distinct metabolic capabilities. Besides, all the top-ranked communities displayed either commensal, mutualistic, or neutral interaction behaviours in the three different nutrient conditions. This suggests that metabolic complementarity and compatibility between the organisms are necessary for the stability of a community.

### Elimination of reactions from competing pathways provide an enhanced lactate flux in the LAB community

Based on the FSEOF (Flux Scanning based on Enforced Objective Flux) approach (see Materials and Methods), we were able to predict suitable reaction knock-outs in six LAB community models that improved lactate flux in comparison to the flux obtained in the wild-type community. These communities each had one organism from the *Leuconostoc* genus, which are capable of fermenting both glucose and xylose. These community species are heterofermentative, i.e., they are capable of production of mixed organic acids such as ethanol, formate, acetate in addition to lactate. Among the predicted knock-out targets, the reactions with a maximum increase of lactate flux are tabulated in Table 1.

**Table 1.** List of reaction knock-outs that lead to increased lactate flux in different LAB communities

| Reaction ID | Reaction Name | Reaction Formula |
| --- | --- | --- |
| **ACKr** | acetate kinase | acetate + ATP ⇌ acetyl-phosphate + ADP |
| **PTAr** | phosphotransacetylase | acetyl-CoA + phosphate ⇌ acetyl-phosphate + CoA |
| **PFL** | pyruvate formate lyase | pyruvate + CoA ⇌ acetyl-CoA + formate |
| **FRD** | fumarate reductase | fumarate + ubiquinol-8 ⇌ succinate + ubiquinone-8 |
| **RPE** | ribulose 5-phosphate 3-epimerase | ribulose 5-phosphate ⇌ xylulose 5-phosphate |
| **XU5PG3PL** | D-xylulose 5-phosphate<br>D-glyceraldehyde-3-phosphate-lyase | xylulose 5-phosphate + phosphate → acetyl-phosphate +<br>glyceraldehyde 3-phosphate + H<sub>2</sub>O |

As evident from these reactions, routes towards the production of other acids, such as acetate, formate, and succinate, are impeded to allow higher flux towards reactions leading to the biosynthesis of lactate. Supplementary Table S4 provides the details of predicted reaction knock-outs in each community model, and the equivalent lactate flux observed in that community upon deletion.

Our findings using this approach for microbial communities concur with experiments observed in literature where deletion of the genes counterpart to these reactions has increased the lactate yield from monocultures of various micro-organisms. An engineered strain of *Enterobacter aerogenes* ATCC 29007 with the phosphate acetyltransferase (*pta*) gene deletion was found to have a higher L-lactate yield by utilization of mannitol (20). *Escherichia coli* K12 strain MG1655 has been engineered by the inactivation of the pyruvate-formate lyase (*pflB*) and fumarate reductase (*frdA*) gene to increase the yield of D-lactate from glycerol (21). A single-gene knock-out of the *pflA* gene in the *E. coli* BW25113 strain has proven to improve D-lactate production from glucose (22). In *Saccharomyces cerevisiae*, the deletion of D-ribulose-5-phosphate 3-epimerase (RPE1) induces the simultaneous utilization of xylose and glucose (23). Gene knock-outs are one of the essential metabolic engineering strategies employed for overcoming barriers of carbon catabolite repression for the co-utilization of carbon sources by microbes (24, 25). Therefore, we hypothesise that to design efficient microbial communities, appropriate gene knock-outs from either one or both the organisms in a co-culture will enhance the co-utilization of mixed carbon substrates. In this regard, in silico approaches as described above will aid in making informed decisions for knock-out experiments.

## Discussion

Lactate synthesis through bacterial fermentation methods is of great importance for improving the compound’s availability and aiding the production of lactate derivatives with high industrial value. While several computational approaches to study microbial communities have emerged in the recent years (6, 26–28), there is still no rigorous methodology to systematically choose a co-culture for optimal production of industrially relevant metabolites, such as the production of lactate. In this study, we report CAMP (Co-culture/Community Analyses for Metabolite Production), an approach to systematically screen multiple candidate communities on multiple substrates under different growth conditions and rank the best performing communities that are most likely to succeed in laboratory experiments. Our approach utilises emerging computational methods with GSMMs in the context of microbial communities of LAB. In pursuit of an ideal two-species community for lactate production, we established a framework where community growth is the objective, and the community model is tested for growth on two primary carbon sources, glucose, and xylose. Screening of viable communities based on predicted growth and lactate yield further enabled comparison between monoculture and co-culture states. Communities were labelled with specific interaction behaviours because of the changes observed in growth rates. The results obtained elucidated the role of single or multi-substrates for the prevalence of a particular interaction type in the communities. Certain cross-fed metabolites among the viable communities were either positively correlated or negatively correlated. This correlation occurred regardless of the interaction type of the community. A change in nutrient condition revealed differences in the interaction behaviours of the communities, but this did not influence the results of the top-ranked communities based on lactate flux. A community comprising of *L. casei* ATCC 334 and *L. plantarum* WCFS1 was selected as the best-performing pair. These species have been used independently in industrial applications as starter cultures. *L. plantarum* is found in many ecological niches and is one of the model organisms in LAB research (29). The GEM of *L. plantarum* was one of the first reported GSMMs from the LAB species (30). The presence of *L. plantarum* in the top-ranked pairs in our study reiterates the compatibility of this microbe with other LAB species and its utility for lactate production. Other *L. plantarum* and *Leuconostoc* species are used as co-cultures for fermentation of Chinese sauerkraut (31). *L. rhamnosus* strains have been co-cultured with *Saccharomyces cerevisiae* for enhanced exopolysaccharide production (32). *Pediococcus acidilactici* species have been co-cultured with *L. delbrueckii* species for pediocin production in milk (33).

Highly efficient micro-organisms are required to meet the industrial standards for lactic acid production. This can be achieved through perturbation, i.e., addition or deletion of genes that enhance the capability of the community to produce lactate. To address this aspect, we undertook an *in silico* strain optimisation approach using FSEOF to predict reactions that can be deleted to improve product flux. The results we observed were encouraging as they were in accordance with previously published experiments where gene deletion was utilised to enhance lactate yield in monocultures of different micro-organisms. These results also allude that gene knock-outs identified in monoculture can be extended to microbial communities as well. The gene knock-outs can be from one or both organisms in a co-culture. Co-cultures and communities of LAB can provide a significant advantage over the engineering of monocultures. With our framework, we have predicted LAB communities, which are useful candidates to produce lactate. These predictions form a ready shortlist for experimental validation. Our workflow can be extended to communities of larger sizes as well, although the increase in combinatorial complexity will also demand an increase in computational cost. The caveat of this study is the dependence on the quality of the GSMMs used. The biochemical pathways to produce the metabolite of interest should also be well defined in the GSMMs. Nevertheless, as newer, more accurate reconstructions emerge, they can be used in our approach to present more accurate insights into the compatibility and interactions between organisms to choose the best possible community for a given application. Our approach provides a ready framework for the integration of additional experimental data arising from transcriptomics studies or 13C metabolic flux analyses, to better constrain the models and improve the accuracy of the predictions.

In sum, we have presented a systematic workflow for the careful screening and analysis of many microbial co-cultures to produce the desired metabolite. Our method examines these co-cultures across growth conditions and across multiple substrates to identify the most promising candidates for experimental validation. Computational approaches, as presented in this study, can provide additional flexibility and valuable insights towards informing the selection of microbial co-cultures for metabolic engineering.

## Materials and Methods

### GSMMs

The Virtual Metabolic Human (www.vmh.life) repository was used for retrieving 47 Lactic Acid Bacteria GSMMs. Models (AGORA version 1.03) of *Lactobacillus, Leuconostoc*, and *Pediococcus* species were obtained (34). Previously curated and published GSMMs of *L. plantarum* WCSF1 and *L. reuteri* JCM 1112 were also used to construct the synthetic communities of LAB (14, 30). A list of all 49 GSMMs used in this study is tabulated in Table S1. Three models from VMH, namely, *L. amylolyticus, L. crispatus*, and *L. delbrueckii subsp. bulgaricus* ATCC BAA 365 did not have the necessary exchange and transport reactions for glucose. We added glucose exchange and transport reactions to these models, based on evidence from literature suggesting their capability to metabolise glucose (35).

### Creation and growth simulations of two-species communities

We generated all possible pairwise combinations of the 49 species to yield 1176 synthetic LAB communities and simulated them using SteadyCom (6), a constraint-based modelling method for the creation and steady-state flux-balance analysis (FBA) of microbial communities. SteadyCom performs a community FBA by computing the relative abundance of each species with the objective function as maximisation of community growth.

LAB are known to be cultured in laboratories with MRS (deMan, Rogosa, and Sharpe) nutrient media. Analogous growth conditions were simulated *in silico* using nutrient uptake components for LAB models obtained from the KOMODO (Known Media Database) at ModelSEED (36). All known 20 amino acids were included in this nutrient media. Lignocellulose hydrolysate contains glucose and xylose as significant components. Hence, to mimic this substrate composition, we constrained the lower bounds of glucose and xylose exchange reactions in the community compartment (u) of the models.

Due to a lack of species-specific data for glucose and xylose uptakes, we considered three nutrient conditions: a) a minimal nutrient condition with −1 mmol/gDW/h of glucose and xylose each, b) an excess nutrient condition with constraints of −30 and −10 mmol/gDW/h for glucose and xylose, respectively, and c) finally a community-specific nutrient condition, where we identified the glucose and xylose uptake fluxes at half-maximal growth rates of each model. The lower bounds of the amino acid exchange reactions and other essential components required for model growth were considered as −1 or −1000 mmol/gDW/h (37). ATP maintenance constraints for all the LAB models were fixed at 0.36 mmol/gDW/h, as observed in the curated *L. plantarum WCFS1* and *L. reuteri* JCM 1112 GSMMs. The growth simulations were performed in an anoxic environment, as LAB are anaerobic micro-organisms. Steady-state community growth rates, as well as species abundances, were computed. All simulations were performed in MATLAB R2018a (MathWorks Inc., USA) using the COBRA Toolbox v3.0 (38) and IBM ILOG CPLEX 12.8 as the linear programming solver.

### Categorising communities based on interaction type

Communities were categorised into six interaction types, namely, parasitism, amensalism, commensalism, mutualism, neutralism, and competitive, based on a 10% difference in growth rates of the microbe when grown in co-culture compared to when the bacterium is grown separately (17). Mutualism and commensalism have a positive effect on community partners, whereas parasitism, competition, and amensalism evoke a negative response on the growth of either partner.

### Studying variation in lactate fluxes in a community using FVA

We calculated the maximum lactate produced by a community using FVA on viable communities. FVA computes the flux range of every reaction by minimising and maximising the flux through the reactions (39). We considered a community to be viable if each organism in the community had a minimum growth rate of 0.01 h^-1^ or higher (40). While performing FVA, the biomass reaction in each community was constrained to the maximum community growth rate obtained. SteadyComFVA was used to calculate the maximum flux through the lactate exchange reaction in the community compartment (“EX_lac_D(u)”).

### Computing expected vs. observed lactate yield in each community

The ConYE model proposed by Medlock *et al*. (41) for identifying metabolic mechanisms of interactions within gut microbiota was adapted to our study to calculate and compare the expected and observed lactate yield from each LAB community. The ConYE model identifies metabolites for which the consumption or production behaviour is altered in co-culture. Each strain is assumed to produce or consume a fixed quantity of each metabolite. This assumption is tested by comparing the expected behaviour to the observed co-culture data. The null hypothesis states that the metabolite in co-culture is equal to the predicted amount. Rejecting the null hypothesis implies that the co-culture has caused at least one species to significantly alter the metabolism of the metabolite (41).

With the lactate fluxes identified in monoculture conditions, an estimate of the lactate flux produced in co-culture can be made, considering the substrate utilisation by each species in co-culture. This computed expected yield of lactate is compared with the maximum lactate fluxes observed in the community compartment (u) in co-culture.

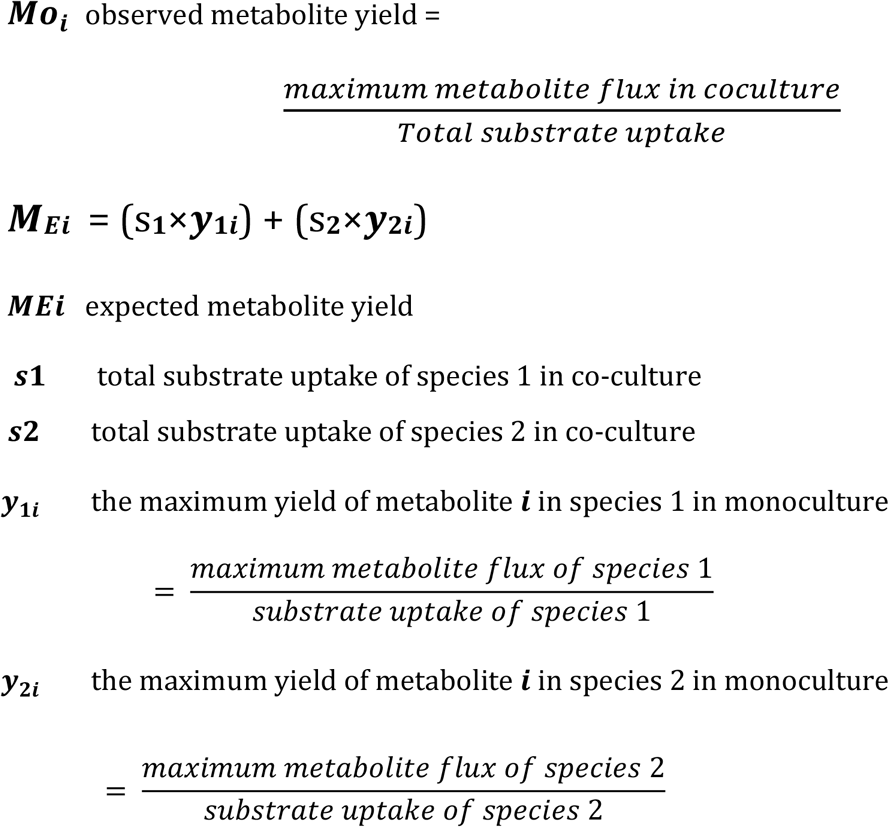

If the observed yield of a community is 10-fold higher than the expected yield, i.e. *M_o_i__* ≥ 10 * *M_E_i’__*, the community is considered as a candidate pair for lactate production.

### Selection of product and growth-efficient communities

Product and growth-efficient communities are defined as communities where a perturbation to the availability of substrates does not affect the viability of the community and the capability to produce lactate. To identify such product and growth-efficient communities, a set of simulations were performed. In the first simulation, the D-Lactate exchange reaction of one organism in the pair was blocked, which prevented cross-feeding of D-Lactate between the community members. Secondly, one organism in the pair was considered as the primary consumer of the substrates, while substrate consumption was blocked in the other organism. Community pairs that retained viability in all simulations were ranked after normalisation (min-max normalisation using the ‘*rescale*’ function in MATLAB R2018a) of lactate yields and growth rates.

### Metabolic Distances of LAB communities

We computed metabolic distances of all LAB models in each community as described in Magnúsdóttir et. al (42). The distance is calculated using the Jaccard distance. Metabolic 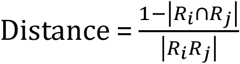, where *R_i_* is the reaction list from the model *i* and *R_j_* is the reaction list of model *j*. Metabolic distance of 1 indicates that the two models do not share any reactions, whereas a metabolic distance of zero indicates that the models have identical reactions.

### Community optimisation and prediction of reaction knock-outs using FSEOF

We performed strain optimisation methods such as the identification of knockout targets in each LAB community that would positively impact lactate production. To this end, we used the FSEOF (Flux Scanning based on Enforced Objective Flux) approach (15). Using FSEOF, potential reactions to be knocked out were selected based on metabolic flux scanning, which selects fluxes towards product formation. Other constraints used to predict reaction knock-outs included an increase in lactate flux of the mutant community model compared to wild-type and viability (i.e., a growth rate of 0.01 h^-1^ or higher) of both organisms in the community. When the number of reactions obtained from FSEOF was less than or equal to an arbitrary threshold of 30, double deletions were carried out to test all possible knock-out combinations (i.e., a maximum of 435 double deletions) of these reactions. The threshold of 30 reactions was chosen for ease of computation. A suitable strategy was selected depending upon the contribution of each deletion towards an increase in lactate flux compared to the wild-type lactate flux. On the other hand, if the reaction list had greater than 30 reactions, only single reaction deletions were performed to identify potential knock-outs that improved lactate flux. For this *in silico* strain optimisation task, the COBRA Toolbox v3.0 functions ‘*removeRxns*’ and ‘*optimizeCbModel*’ were used for reaction deletions and FBA with optimisation of community biomass, respectively.

## Data availability

All models used in this work and the codes used for our analysis are available at: https://github.com/RamanLab/CAMP

## Acknowledgments

M.I. acknowledges the IIT Madras Institute Post-Doctoral Fellowship and the Post-Doctoral fellowship from Initiative for Biological Systems Engineering (IBSE), IIT Madras, India.

---

**Fig. S1:**
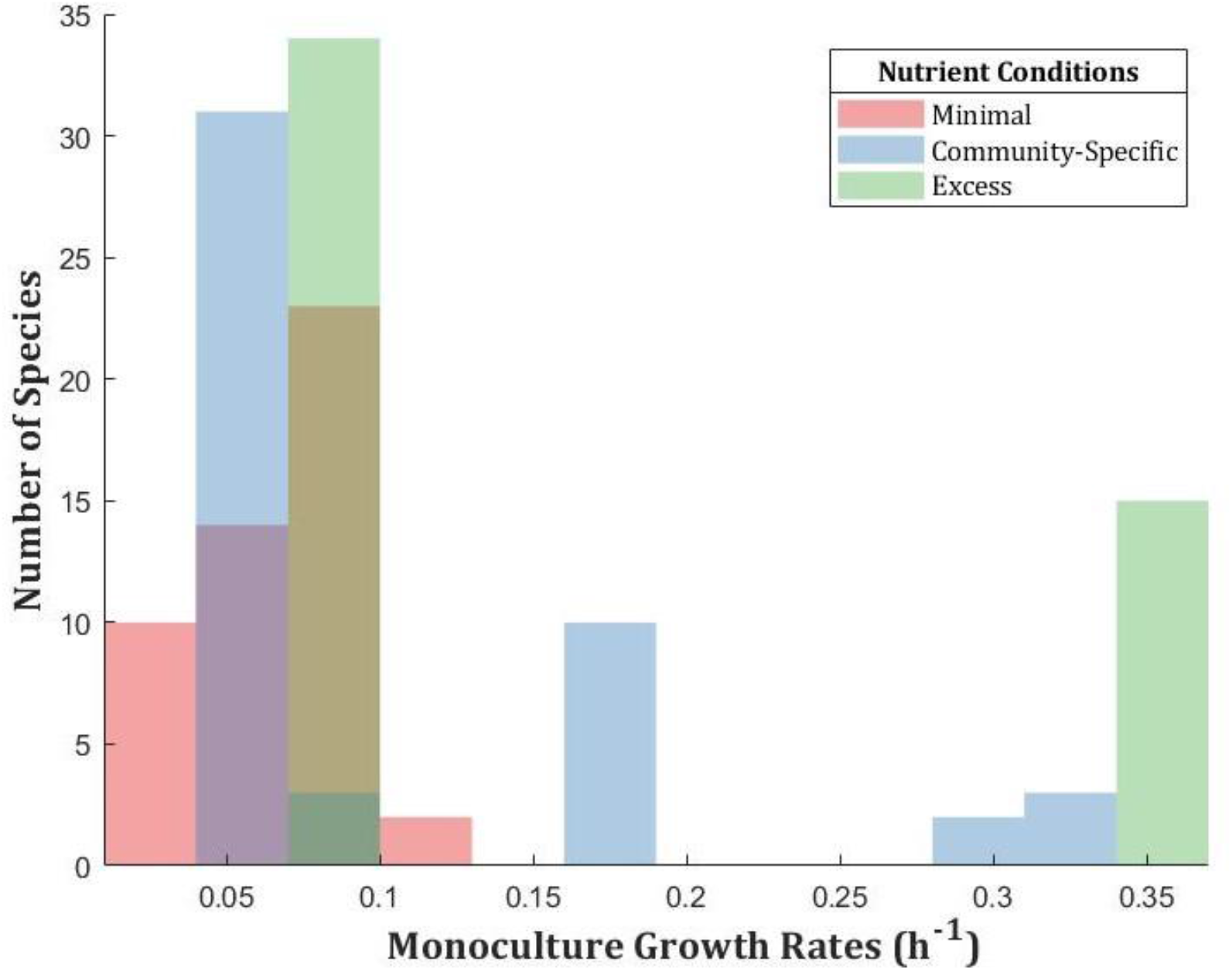
Histogram distribution of monoculture growth rates of all 49 species under three different nutrient conditions

**Fig. S2:**
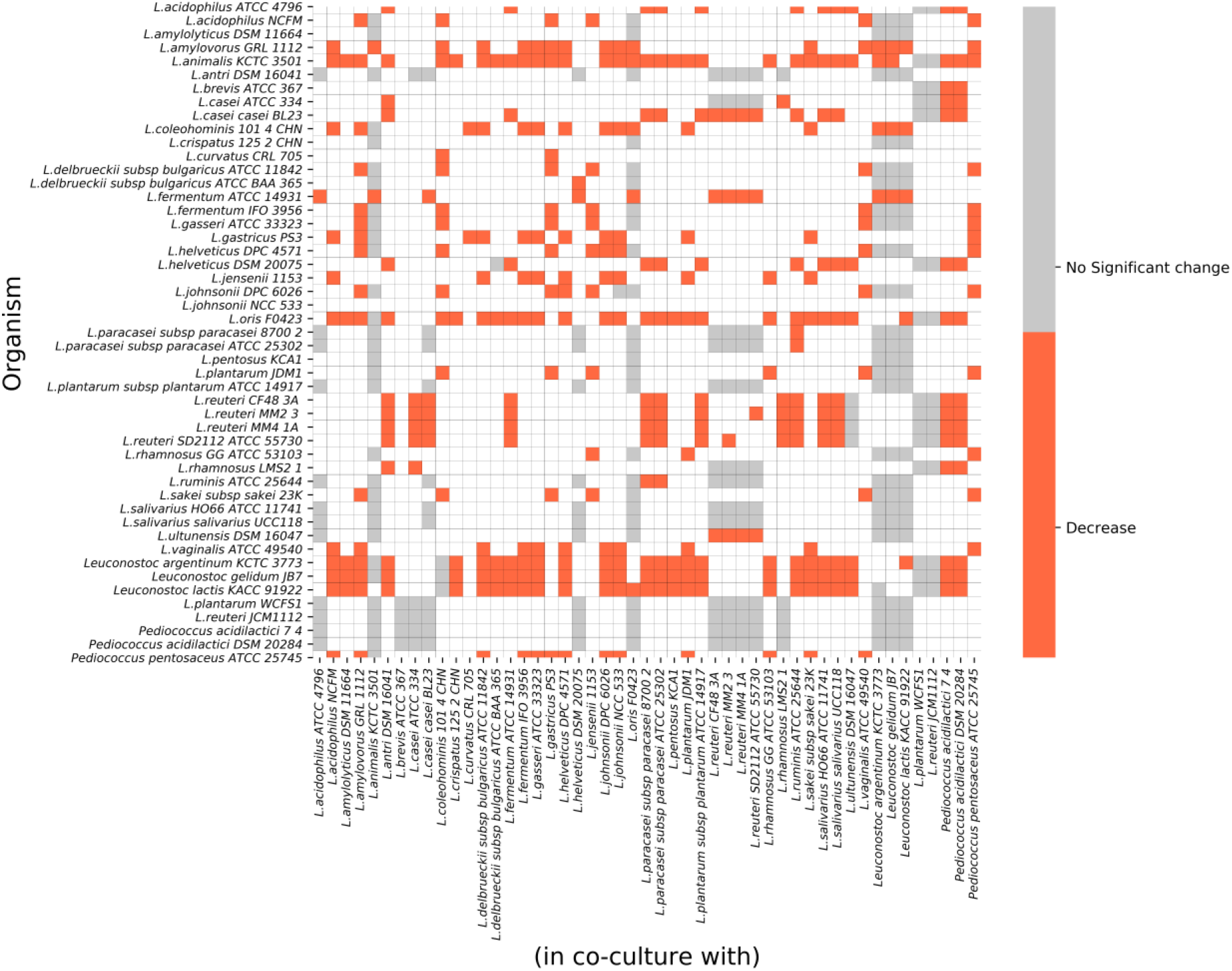
Monoculture vs. Co-culture growth rates with excess nutrient uptake. The heatmap depicts the change in the growth rate of an organism’s monoculture growth compared to when it is co-cultured with another species under excess nutrient uptake condition. A difference lesser than 10% of monoculture growth is regarded as a decrease. 838 non-viable pairs and the diagonal, which represents 49 monocultures, are depicted as white squares.

**Fig. S3:**
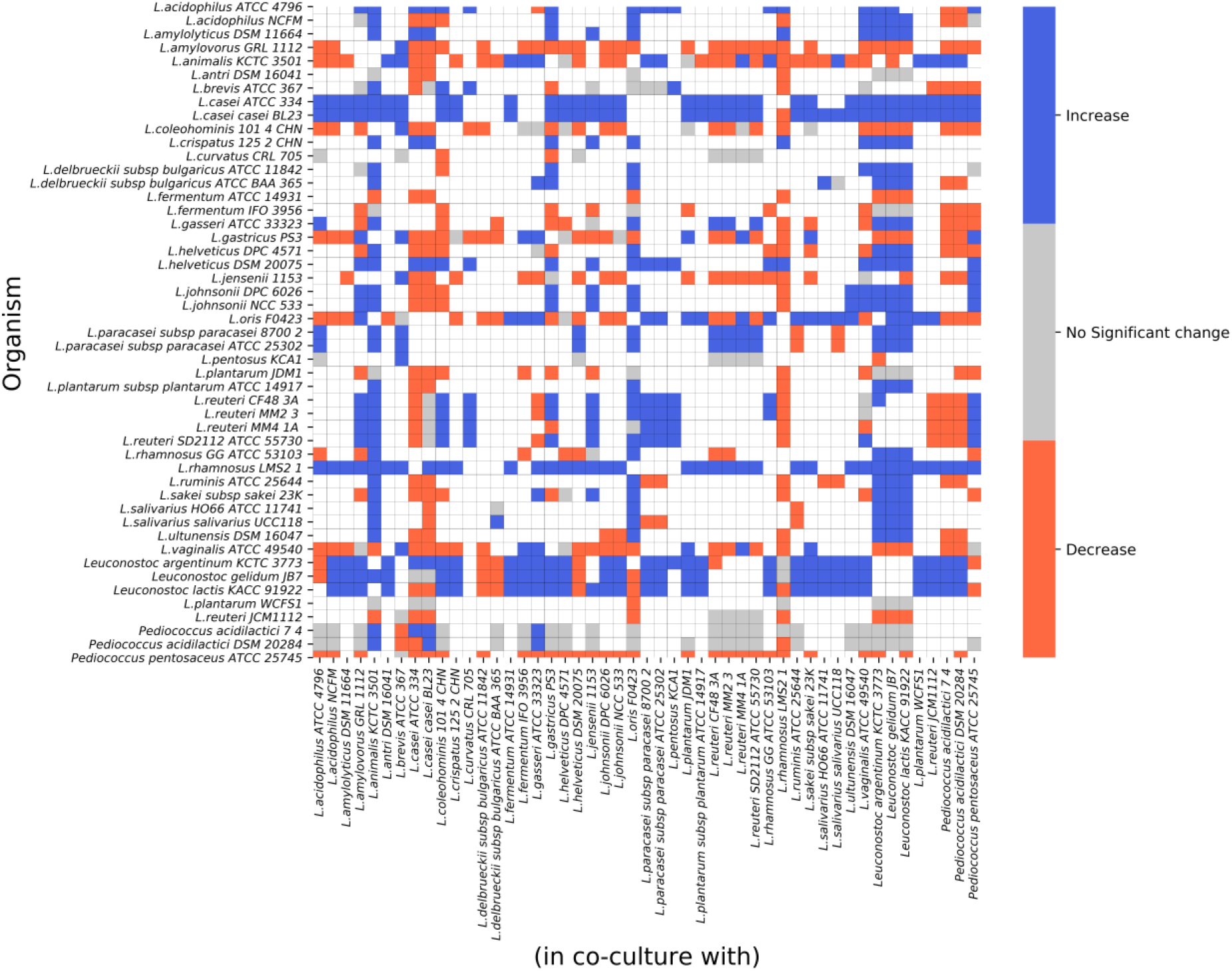
Monoculture vs. Co-culture growth rates with minimal nutrient uptake. The heatmap depicts the change in the growth rate of an organism’s monoculture growth compared to when it is co-cultured with another species under minimal nutrient uptake condition. A difference greater than 10% of monoculture growth is considered an increase, lesser than 10% of monoculture growth is regarded as a decrease. 684 non-viable pairs and the diagonal, which represents 49 monocultures, are depicted as white squares.

**Fig. S4:**
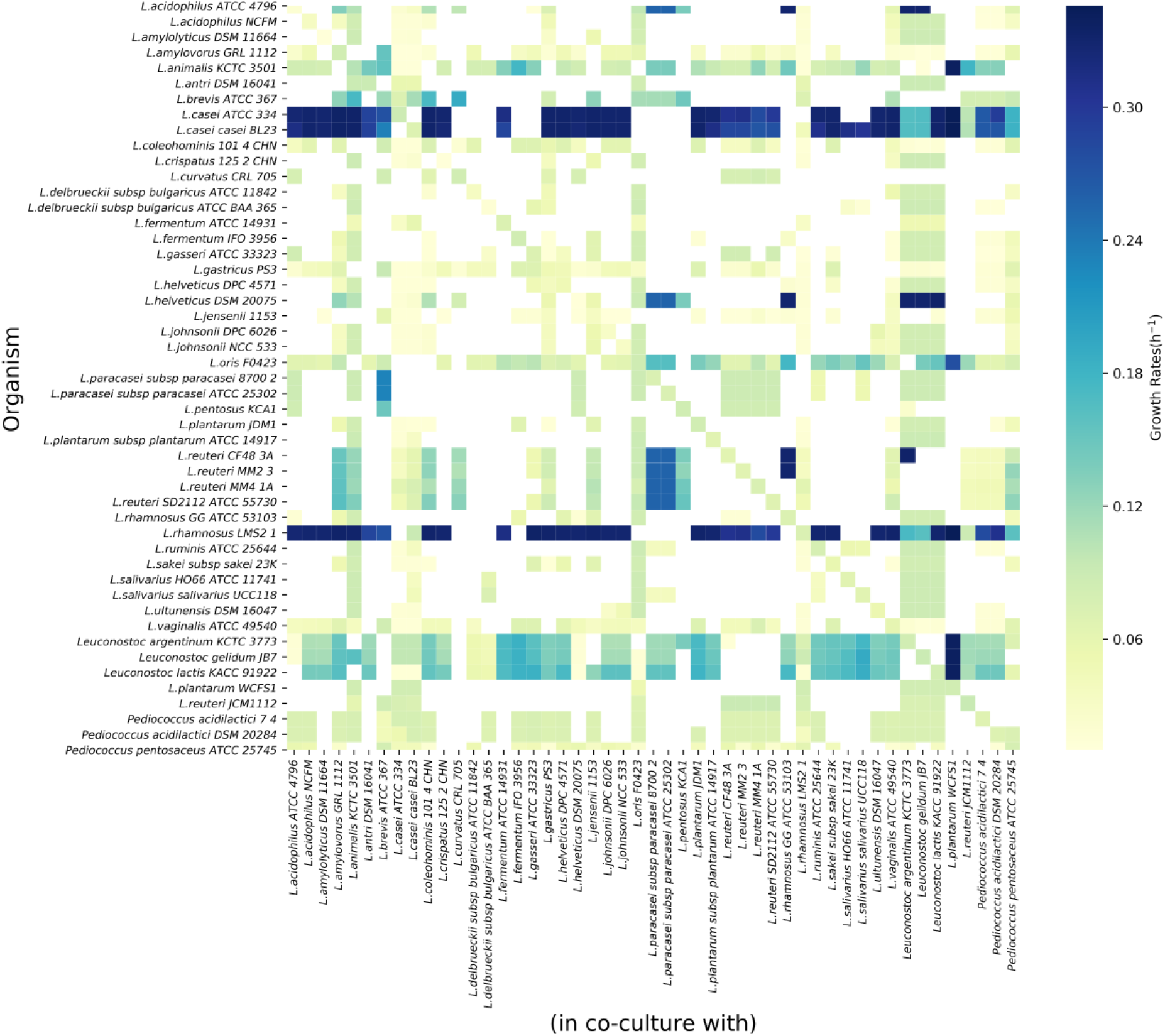
Monoculture and Coculture growth rates with minimal nutrient uptake. The heatmap depicts the absolute values of the predicted growth rates of each organism in the community. Diagonal elements represent the monoculture growth rates of all 49 species. Non-viable communities are denoted in white squares.

**Fig. S5:**
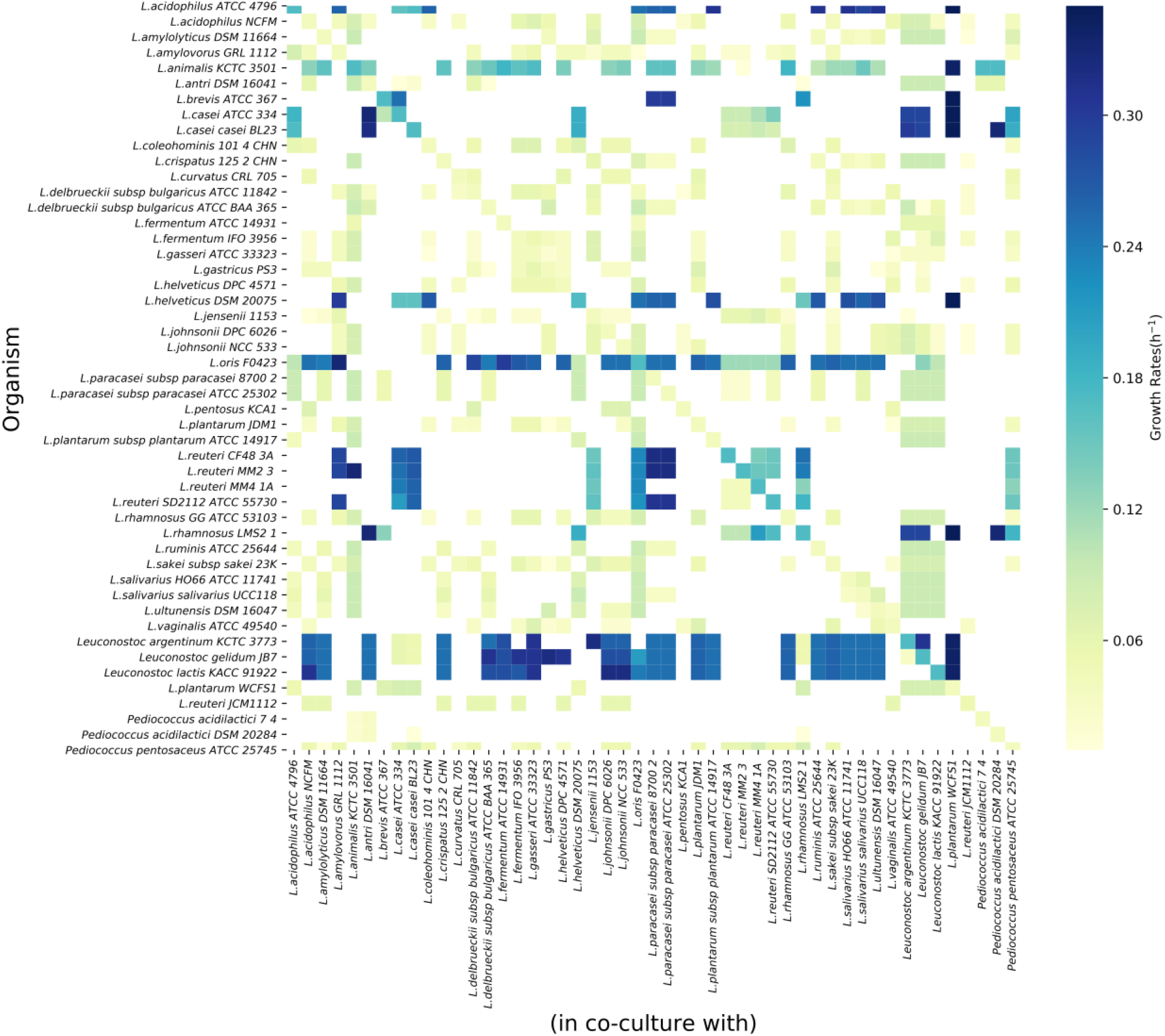
Monoculture and Coculture growth rates with community-specific nutrient uptake fluxes. The heatmap depicts the absolute values of the predicted growth rates of each organism in the community. Diagonal elements represent the monoculture growth rates of all 49 species. Non-viable communities are denoted in white squares.

**Fig. S6:**
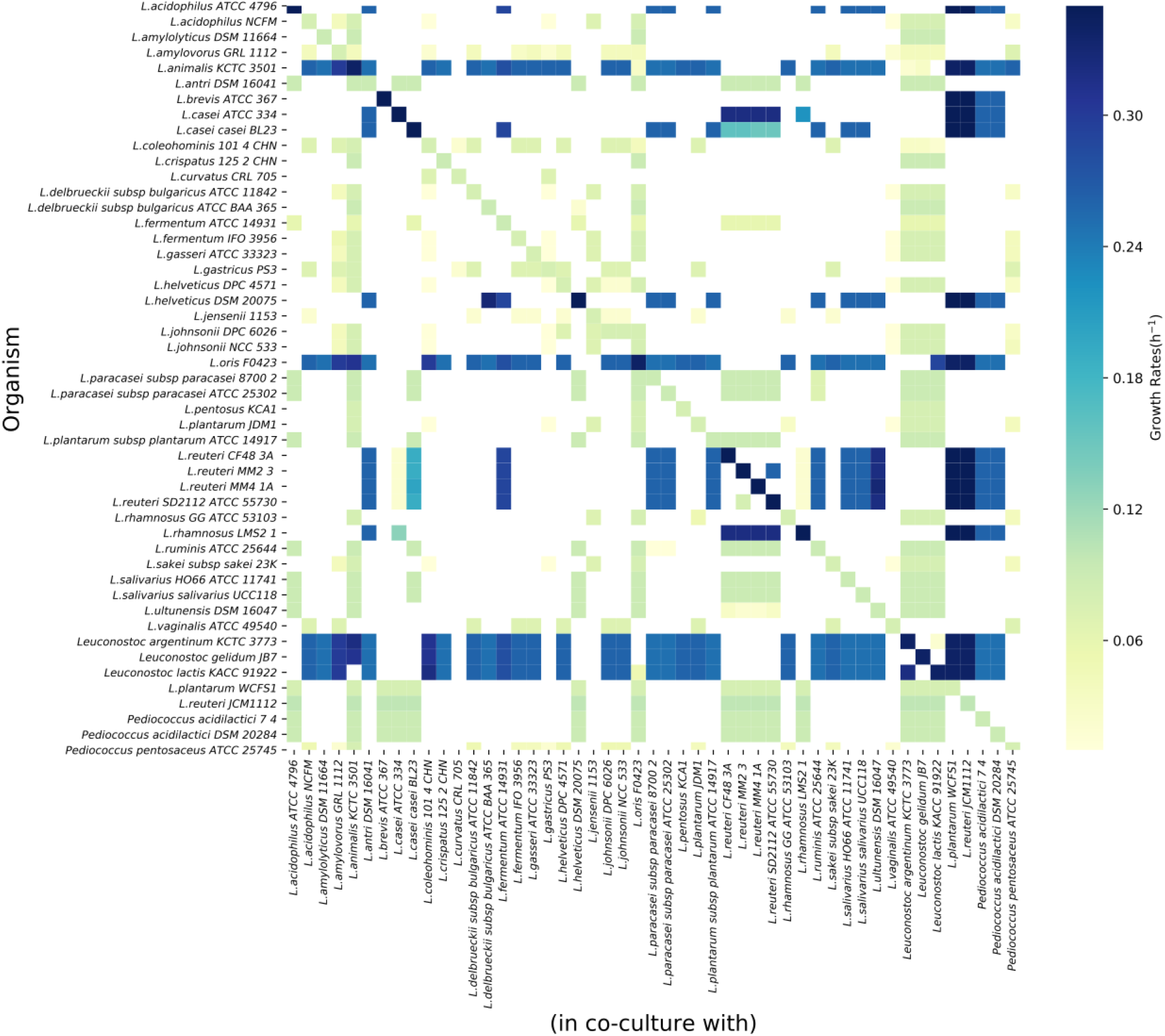
Monoculture and Coculture growth rates in excess-nutrient condition. The heatmap depicts the absolute values of the predicted growth rates of each organism in the community. Diagonal elements represent the monoculture growth rates of all 49 species. Non-viable communities are denoted in white squares.

**Fig. S7:**
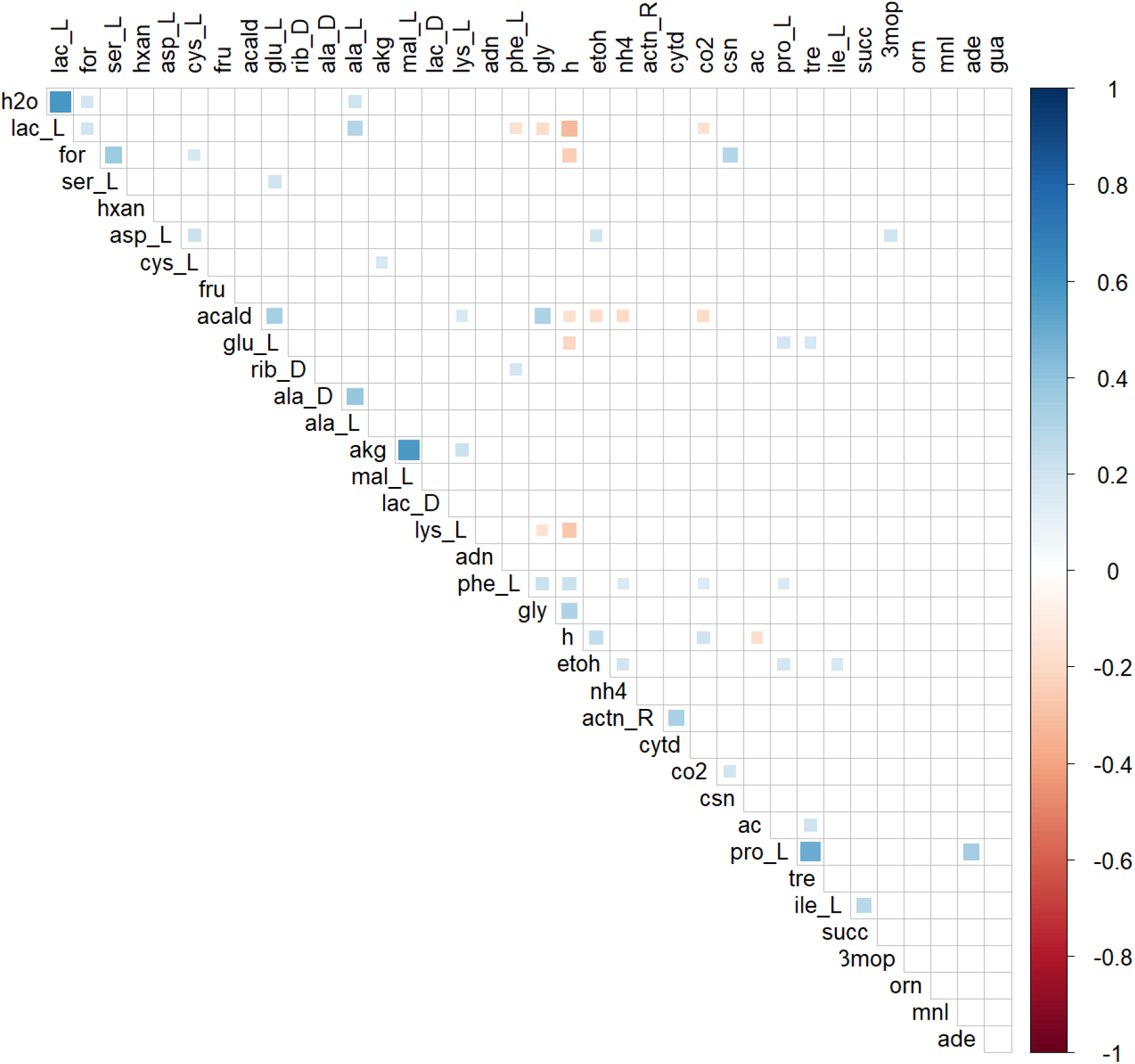
Correlation between the cross-fed metabolites in the excess nutrient condition. Positively correlated metabolites are denoted in blue, whereas negatively correlated metabolites are denoted in brown. Alpha-ketoglutarate and malate, Proline and trehalose are among the positively correlated metabolites.

**Fig. S8:**
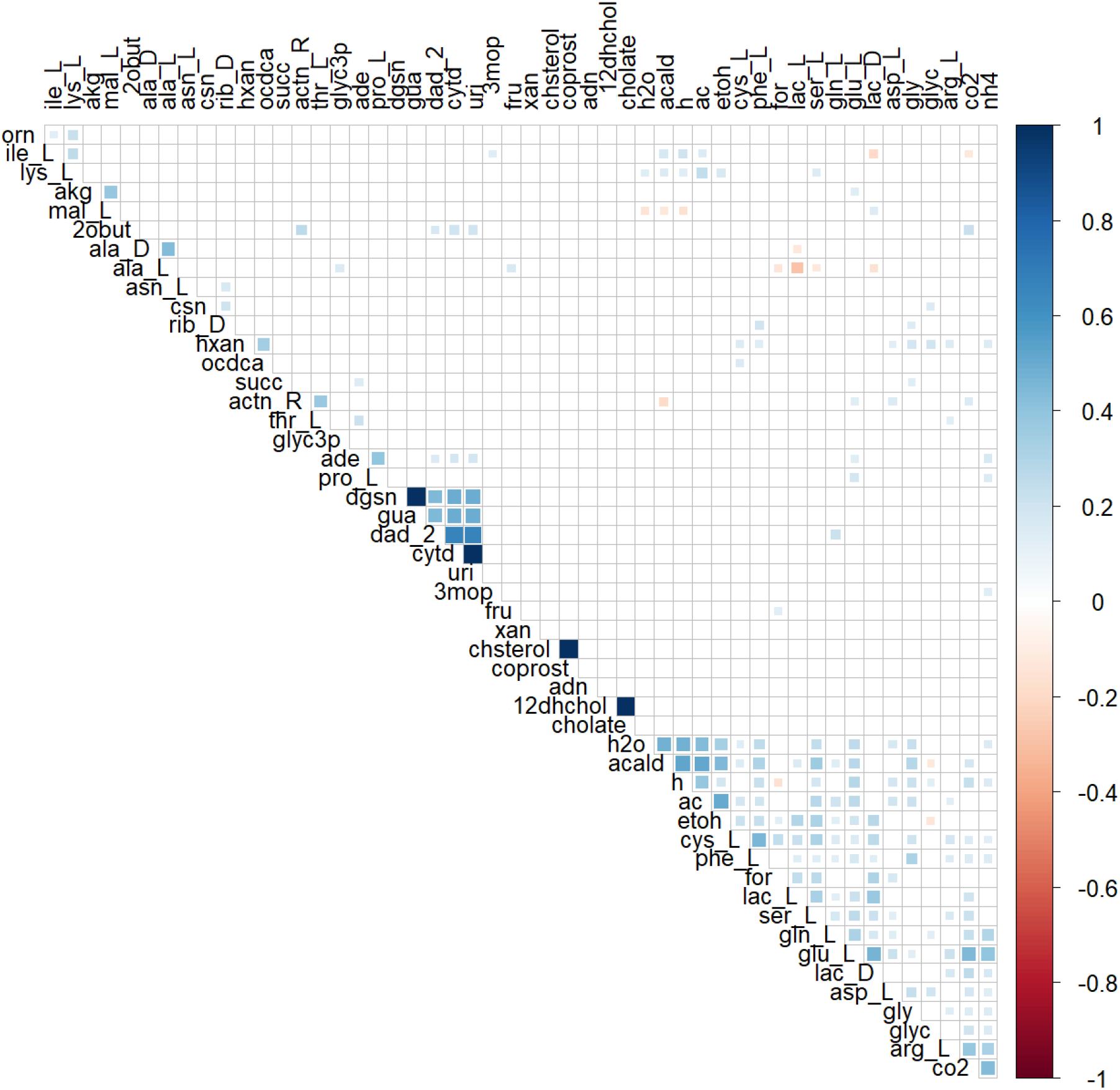
Correlation between the cross-fed metabolites in the minimal nutrient condition. Positively correlated metabolites are denoted in blue, whereas negatively correlated metabolites are denoted in brown. Acetate and acetaldehyde, ethanol and acetaldehyde are among the positively correlated cross-fed metabolites.

## References

1. Eş I, Mousavi Khaneghah A, Barba FJ, Saraiva JA, Sant’Ana AS, Hashemi SMB. 2018. Recent advancements in lactic acid production - a review. Food Res Int 107:763–770.

2. Jawed K, Yazdani SS, Koffas MA. 2019. Advances in the development and application of microbial consortia for metabolic engineering. Metab Eng Commun 9:e00095.

3. Gu C, Kim GB, Kim WJ, Kim HU, Lee SY. 2019. Current status and applications of genome-scale metabolic models. Genome Biol 20:1–18.

4. Stolyar S, Van Dien S, Hillesland KL, Pinel N, Lie TJ, Leigh JA, Stahl DA. 2007. Metabolic modeling of a mutualistic microbial community. Mol Syst Biol 3:1–14.

5. Ye C, Zou W, Xu N, Liu L. 2014. Metabolic model reconstruction and analysis of an artificial microbial ecosystem for vitamin C production. J Biotechnol 182–183:61–67.

6. Chan SHJ, Simons MN, Maranas CD. 2017. SteadyCom: Predicting microbial abundances while ensuring community stability. PLoS Comput Biol 13:1–25.

7. Dusselier M, Van Wouwe P, Dewaele A, Makshina E, Sels BF. 2013. Lactic acid as a platform chemical in the biobased economy: The role of chemocatalysis. Energy Environ Sci 6:1415–1442.

8. Farah S, Anderson DG, Langer R. 2016. Physical and mechanical properties of PLA, and their functions in widespread applications — A comprehensive review. Adv Drug Deliv Rev 107:367–392.

9. Alves de Oliveira R, Komesu A, Vaz Rossell CE, Maciel Filho R. 2018. Challenges and opportunities in lactic acid bioprocess design—From economic to production aspects. Biochem Eng J 133:219–239.

10. Juturu V, Wu JC. 2016. Microbial production of lactic acid: the latest development. Crit Rev Biotechnol 36:967–977.

11. Zhang Y, Vadlani P V. 2015. Lactic acid production from biomass-derived sugars via co-fermentation of Lactobacillus brevis and Lactobacillus plantarum. J Biosci Bioeng 119:694–699.

12. Cui F, Li Y, Wan C. 2011. Lactic acid production from corn stover using mixed cultures of Lactobacillus rhamnosus and Lactobacillus brevis. Bioresour Technol 102:1831–1836.

13. Tarraran L, Mazzoli R. 2018. Alternative strategies for lignocellulose fermentation through lactic acid bacteria: The state of the art and perspectives. FEMS Microbiol Lett 365.

14. Kristjansdottir T, Bosma EF, Branco Dos Santos F, Özdemir E, Herrgård MJ, França L, Ferreira B, Nielsen AT, Gudmundsson S. 2019. A metabolic reconstruction of Lactobacillus reuteri JCM 1112 and analysis of its potential as a cell factory. Microb Cell Fact 18:1–19.

15. Choi HS, Lee SY, Kim TY, Woo HM. 2010. In silico identification of gene amplification targets for improvement of lycopene production. Appl Environ Microbiol 76:3097–3105.

16. Spector MP. 2009. Encyclopedia of Microbiology. Encycl Microbiol 242–264.

17. Heinken A, Thiele I. 2015. Anoxic conditions promote species-specific mutualism between gut microbes In Silico. Appl Environ Microbiol 81:4049–4061.

18. D’Souza G, Shitut S, Preussger D, Yousif G, Waschina S, Kost C. 2018. Ecology and evolution of metabolic cross-feeding interactions in bacteria. Nat Prod Rep 35:455–488.

19. Pacheco AR, Moel M, Segrè D. 2019. Costless metabolic secretions as drivers of interspecies interactions in microbial ecosystems. Nat Commun 10.

20. Thapa LP, Lee SJ, Park C, Kim SW. 2017. Production of L-lactic acid from metabolically engineered strain of Enterobacter aerogenes ATCC 29007. Enzyme Microb Technol 102:1–8.

21. Mazumdar S, Clomburg JM, Gonzalez R. 2010. Escherichia coli strains engineered for homofermentative production of D-lactic acid from glycerol. Appl Environ Microbiol 76:4327–4336.

22. Zhu J, Shimizu K. 2005. Effect of a single-gene knockout on the metabolic regulation in Escherichia coli for D-lactate production under microaerobic condition. Metab Eng 7:104–115.

23. Shen MH, Song H, Li BZ, Yuan YJ. 2015. Deletion of d-ribulose-5-phosphate 3-epimerase (RPE1) induces simultaneous utilization of xylose and glucose in xylose-utilizing Saccharomyces cerevisiae. Biotechnol Lett 37:1031–1036.

24. Wu Y, Shen X, Yuan Q, Yan Y. 2016. Metabolic engineering strategies for coutilization of carbon sources in microbes. Bioengineering 3:1–10.

25. Chiang C, Lee HM, Guo HJ, Wang ZW, Lin L. 2013. Systematic Approach To Engineer Escherichia coli Pathways for.

26. Khandelwal RA, Olivier BG, Röling WFM, Teusink B, Bruggeman FJ. 2013. Community Flux Balance Analysis for Microbial Consortia at Balanced Growth. PLoS One 8.

27. Zomorrodi AR, Maranas CD. 2012. OptCom: A multi-level optimization framework for the metabolic modeling and analysis of microbial communities. PLoS Comput Biol 8.

28. Ravikrishnan A, Raman K. 2018. Systems-Level Modelling of Microbial Communities1st Editio. CRC Press.

29. Siezen RJ, van Hylckama Vlieg JET. 2011. Genomic diversity and versatility of Lactobacillus plantarum, a natural metabolic engineer. Microb Cell Fact 10:1–13.

30. Teusink B, Wiersma A, Molenaar D, Francke C, De Vos WM, Siezen RJ, Smid EJ. 2006. Analysis of growth of Lactobacillus plantarum WCFS1 on a complex medium using a genome-scale metabolic model. J Biol Chem 281:40041–40048.

31. Xiong T, Peng F, Liu Y, Deng Y, Wang X, Xie M. 2014. Fermentation of Chinese sauerkraut in pure culture and binary co-culture with Leuconostoc mesenteroides and Lactobacillus plantarum. LWT - Food Sci Technol 59:713–717.

32. Bertsch A, Roy D, LaPointe G. 2019. Enhanced exopolysaccharide production by Lactobacillus rhamnosus in co-culture with Saccharomyces cerevisiae. Appl Sci 9.

33. Somkuti GA, Steinberg DH. 2010. Pediocin production in milk by Pediococcus acidilactici in co-culture with Streptococcus thermophilus and Lactobacillus delbrueckii subsp. bulgaricus. J Ind Microbiol Biotechnol 37:65–69.

34. Noronha A, Modamio J, Jarosz Y, Guerard E, Sompairac N, Preciat G, Daníelsdóttir AD, Krecke M, Merten D, Haraldsdóttir HS, Heinken A, Heirendt L, Magnúsdóttir S, Ravcheev DA, Sahoo S, Gawron P, Friscioni L, Garcia B, Prendergast M, Puente A, Rodrigues M, Roy A, Rouquaya M, Wiltgen L, Žagare A, John E, Krueger M, Kuperstein I, Zinovyev A, Schneider R, Fleming RMT, Thiele I. 2019. The Virtual Metabolic Human database: Integrating human and gut microbiome metabolism with nutrition and disease. Nucleic Acids Res 47:D614–D624.

35. Carr FJ, Chill D, Maida N. 2002. The lactic acid bacteria: A literature survey. Crit Rev Microbiol 28:281–370.

36. Henry CS, Dejongh M, Best AA, Frybarger PM, Linsay B, Stevens RL. 2010. High-throughput generation, optimization and analysis of genome-scale metabolic models. Nat Biotechnol 28:977–982.

37. Bauer E, Thiele I. 2018. From metagenomic data to personalized in silico microbiotas: predicting dietary supplements for Crohn’s disease. npj Syst Biol Appl 4.

38. Heirendt L, Arreckx S, Pfau T, Mendoza SN, Richelle A, Heinken A, Haraldsdóttir HS, Wachowiak J, Keating SM, Vlasov V, Magnusdóttir S, Ng CY, Preciat G, Žagare A, Chan SHJ, Aurich MK, Clancy CM, Modamio J, Sauls JT, Noronha A, Bordbar A, Cousins B, El Assal DC, Valcarcel L V., Apaolaza I, Ghaderi S, Ahookhosh M, Ben Guebila M, Kostromins A, Sompairac N, Le HM, Ma D, Sun Y, Wang L, Yurkovich JT, Oliveira MAP, Vuong PT, El Assal LP, Kuperstein I, Zinovyev A, Hinton HS, Bryant WA, Aragón Artacho FJ, Planes FJ, Stalidzans E, Maass A, Vempala S, Hucka M, Saunders MA, Maranas CD, Lewis NE, Sauter T, Palsson B, Thiele I, Fleming RMT. 2019. Creation and analysis of biochemical constraint-based models using the COBRA Toolbox v.3.0. Nat Protoc 14:639–702.

39. Mahadevan R, Schilling CH. 2003. The effects of alternate optimal solutions in constraint-based genome-scale metabolic models. Metab Eng 5:264–276.

40. Devika NT, Raman K. 2019. Deciphering the metabolic capabilities of Bifidobacteria using genome-scale metabolic models. Sci Rep 9:1–9.

41. Medlock GL, Carey MA, McDuffie DG, Mundy MB, Giallourou N, Swann JR, Kolling GL, Papin JA. 2018. Inferring Metabolic Mechanisms of Interaction within a Defined Gut Microbiota. Cell Syst 7:245–257.e7.

42. Magnúsdóttir S, Heinken A, Kutt L, Ravcheev DA, Bauer E, Noronha A, Greenhalgh K, Jäger C, Baginska J, Wilmes P, Fleming RMT, Thiele I. 2017. Generation of genome-scale metabolic reconstructions for 773 members of the human gut microbiota. Nat Biotechnol 35:81–89.

